# Decrypting corals: Does regulatory evolution underlie environmental specialization of coral cryptic lineages?

**DOI:** 10.1101/2023.10.11.561896

**Authors:** Dominique N Gallery, John P Rippe, Mikhail V Matz

**Affiliations:** Department of Integrative Biology, University of Texas at Austin, Austin, TX, USA

**Keywords:** gene expression divergence, cryptic speciation, ecological specialization, local adaptation

## Abstract

A recent study has shown that two common Caribbean corals, *Montastrea cavernosa* and *Siderastrea siderea*, in the Florida Keys each consist of four genetically distinct lineages. These lineages are strongly specialized to a certain depth and, to a lesser extent, to nearshore or offshore habitat. We hypothesized that the lineages’ environmental specialization is at least in part due to regulatory evolution, which would be manifested as the emergence of groups of coregulated genes (“modules”) demonstrating lineage-specific responses to different reef environments. Our hypothesis also predicted that genes belonging to such modules would show greater genetic divergence between lineages than other genes. Contrary to these expectations, responses of cryptic lineages to natural environmental variation were essentially the same at the genome-wide gene coexpression network level. Moreover, none of the identified coregulated gene modules exhibit elevated between-lineage divergence. The environmental specialization of cryptic lineages must, therefore, come from relatively subtle adjustments to gene function or regulation that are not detectable at the gene network level and/or involve other constituents of the coral holobiont rather than the coral host.

## Introduction

As anthropogenic stressors increase, coral populations will continue to decline, reducing genetic diversity and, thus, corals’ ability to adapt to future conditions (Carpenter et al., 2008). The challenges of managing coral genetic diversity are complicated by increasing evidence for highly distinct genetic lineages within nominal species (Hays et al., 2021; Knowlton, 1993). Limited introgression between these lineages exacerbates genetic diversity loss due to restricted gene flow (Bálint et al., 2011) and makes it challenging for managers to make informed conservation decisions during restoration. While such lineages are known from many coral species (Afiq-Rosli et al., 2021; Gijsbers et al., 2023; Gómez-Corrales & Prada, 2020; Johnston et al., 2021; Ladner & Palumbi, 2012; Nakajima et al., 2012; Oku et al., 2020; Pinzón & Weil, 2011; Pipithkul et al., 2021; Rippe et al., 2021; Rosser, 2015; Sheets et al., 2018; Stefani et al., 2011; Zayasu et al., 2021), we currently lack an understanding of how these lineages arise and what prevents them from merging.

Current hypotheses suggest that one of the main forces responsible for the lineages’ existence is strong spatially varying selection reinforcing their environmental specialization (Prada & Hellberg, 2021; Sanford & Kelly, 2011). Examining their gene expression is one way to investigate how different lineages adapt or acclimatize to their habitat. Here, we studied two ubiquitous Caribbean species with known cryptic genetic diversity, *Montastraea cavernosa* and *Siderastrea siderea*. These annually gonochoristic species are broadcast spawning species with larval durations of approximately two weeks (Frys et al., 2020). In the Florida Keys, these species are found across the reef shelf encompassing nearshore shallow, offshore shallow, and offshore deep habitats (Figure 1a) with cryptic genetic lineages specialized to either the shallow or deep habitats (Rippe et al., 2021). These habitats vary in their environmental parameters, with the nearshore shallow habitat exhibiting larger temperature fluctuations, increased turbidity, and higher terrestrial nutrients (Lirman & Fong, 2007). The offshore shallow habitat exhibits reduced temperature variability and a reduction of terrestrial impacts (e.g., turbidity and nutrients) (Kenkel & Matz, 2016; Lapointe et al., 2004; Lee & Williams, 1999), whereas the offshore deep has the least light of the three sites, and also differs in inorganic nutrients (*2020 FKNMS Annual Report.Pdf*, n.d.).

**Figure 1:**
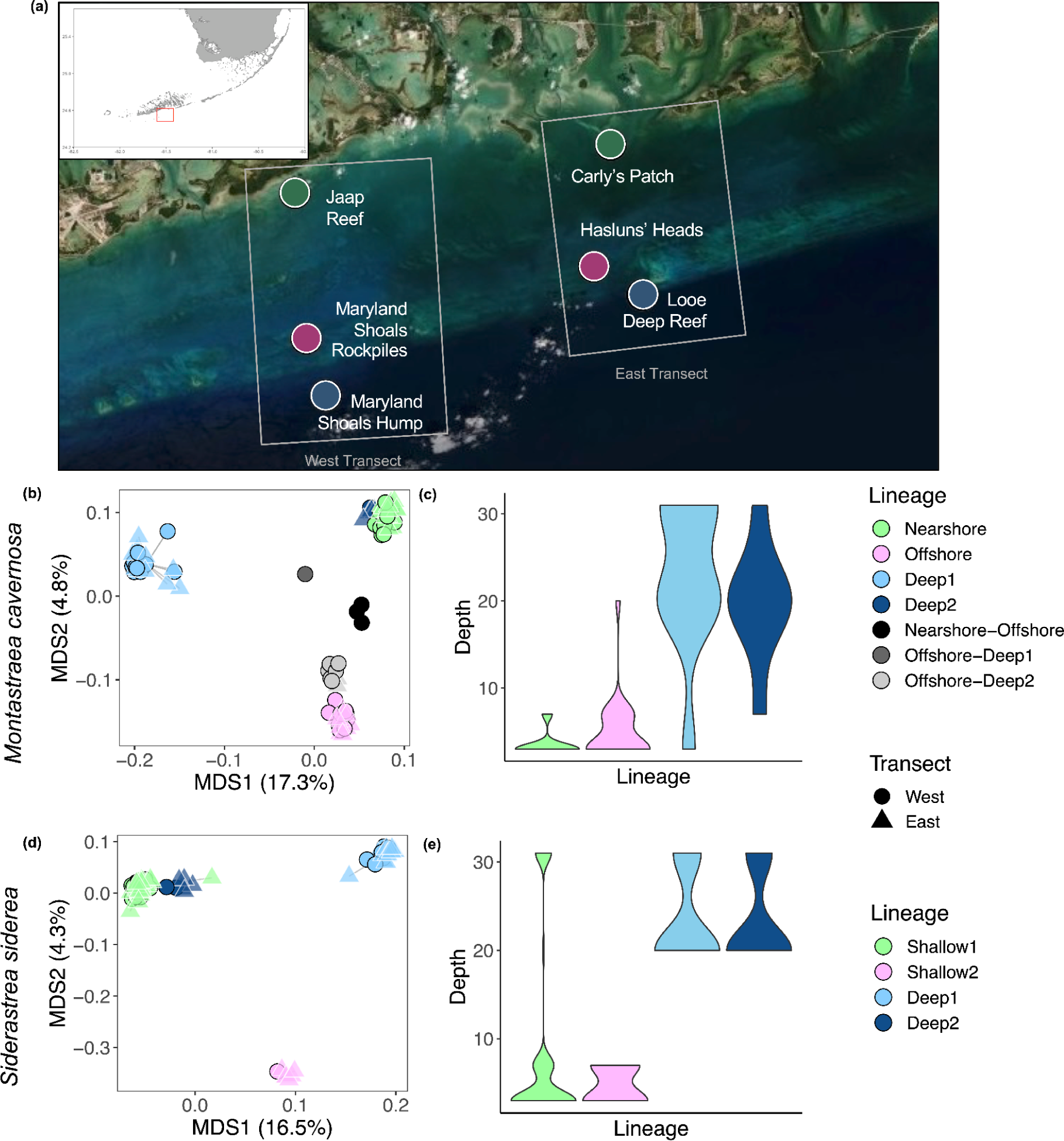
Distribution of cryptic lineages across cross-shelf gradient. (a) Sampling sites: the East transect (Rippe et al. 2021) and the West transect (this study). Cryptic genetic structure (b) and depth distribution of lineages (c) of *Montastrea cavernosa*. Panels (d-e) show the same for *Siderastrea siderea*. (b,d) Principal coordinate analyses of both transects. West transect individuals are indicated by a circle, and east transect individuals are indicated by a triangle. (c, e) Distribution of each lineage across depths (in meters) for non-hybrid individuals of *M. cavernosa* (c) and *S. siderea* (e) for both transects. Individuals with greater than 25% identification with more than one lineage were considered hybrids. Lineages are named as in Rippe et al. (2021).

Here, we confirmed the uneven distribution of these lineages along an independent cross-shelf transect and investigated the underlying mechanisms of their differential adaptation by looking at their gene expression in the wild. We expected to see lineage-dependent gene expression responses to the three habitat types (nearshore, offshore, deep), reflecting their differential adaptation to these habitats. Furthermore, we predicted that genes constituting these lineage-specific responses would show more genetic divergence between lineages compared to the rest of the genes in the genome. Together, these observations would indicate that regulatory evolution has contributed to the ecological specialization of the cryptic lineages.

## Methods

### Sample collection

*Montastraea cavernosa* and *Siderastrea siderea* tissue samples were collected from colonies from three habitats in the lower Florida Keys: nearshore shallow, offshore shallow, and offshore deep (Figure 1a). Approximately 20 colonies >30 cm in diameter were collected for each species from three sites: Jaap Reef (24.5857°N, 81.58262°W; nearshore shallow) at 3m, Maryland Shoals Rockpiles (24.52172°N, 81.57675°W; offshore shallow) at 3m, and Maryland Shoals Hump (24.49659°N, 81.56746°W; offshore deep) at 31m (west transect, Figure 1a). Following collection, samples were transported to shore on ice, preserved in 100% ethanol at −20°C at Mote Marine’s Elizabeth Moore International Center for Coral Reef Research and Restoration field station in Summerland Key. Once transported to the University of Texas at Austin, samples were stored at −80°C until processed.

### Library preparation and sequencing

DNA and RNA were extracted from tissue samples using RNAqueous Total RNA Isolation Kit (Thermo Fisher Scientific, AM1912) following the manufacturer’s protocol. Eluted samples were divided into two vials for separate RNA and DNA processing and treated with DNase or RNase, respectively. RNA processing for TagSeq sequencing (Meyer et al., 2011) followed the current protocol available at https://github.com/z0on/tag-based_RNAseq. Briefly, ∼2µg of RNA was treated with 1.5µL of DNase prior to RNA fragmentation. cDNA was synthesized using SMARTScribe Reverse Transcriptase (Takara, CAT# 639537) and PCR-amplified 2.5 mM DNTPs (NEB, CAT#N0447S), 10X PCR buffer (Klentaq, CAT#RB20), Klentaq DNA polymerase (Klentaq, CAT#100), and 10 µM 5ILL oligo (IDT). Each sample was uniquely barcoded and sequenced on the Illumina HiSeq 2500 platform at the University of Texas at Austin Genomic Sequencing and Analysis Facility.

DNA processing followed the same methods as Rippe et al. (2021). DNA samples were cleaned with Genomic DNA Clean & Concentrator-10 (Zymo, D4011) prior to preparation for 2bRAD sequencing (Wang et al., 2012). The protocol for laboratory preparation of 2bRAD sequencing is available at https://github.com/z0on/2bRAD_denovo. Sequencing was performed on the Illumina HiSeq 2500 platform at the University of Texas at Austin Genomic Sequencing and Analysis Facility.

### Population structure and confirmation of lineage

To determine if the lineages previously described in Rippe et al. (2021) were present at the second transect, 2bRAD raw sequences were processed following the same methods described in Rippe et al. (2021). Briefly, *M. cavernosa* reads were cleaned and demultiplexed following the referenced-based walkthrough (https://github.com/z0on/2bRAD_denovo/blob/master/2bRAD_README.sh), quality filtered with CUTADAPT version 3.5 (Martin, 2011), and mapped to the *M. cavernosa* genome (Rippe et al., 2021); https://www.dropbox.com/s/yfqefzntt896xfz/Mcavernosa_genome.tgz) with bowtie2 version 2.4.5 (Langmead & Salzberg, 2012). *Siderastrea siderea* reads were cleaned and demultiplexed following the *de novo*walkthrough (https://github.com/z0on/2bRAD_denovo/blob/master/2bRAD_README.sh). Because there is no genome for *S. siderea*, reads were first mapped to concatenated Symbiodiniaceae genomes (Dougan et al., 2022; Scott et al., 2022; Shoguchi et al., 2013, 2018) with bowtie2 version 2.4.5 (Langmead & Salzberg, 2012). Any mapped reads were discarded to remove symbiont contamination from coral host reads. Coral host reads were combined with those of the first transect (Rippe et al., 2021) to create the cluster-derived reference (Wang et al., 2012) following steps outlined at https://github.com/z0on/2bRAD_denovo/) Samples from the second transect were then mapped to this reference using bowtie2 version 2.4.5 (Langmead & Salzberg, 2012).

Following mapping, files were converted to BAM format using SAMtools version 1.15 (Danecek et al., 2021; Li et al., 2009). Sample quality and coverage depth were calculated using ANGSD version 0.937 (Korneliussen et al., 2014), and samples with less than 5% mapping efficiency were removed from downstream analyses (*M. cavernosa* n=5, *S. siderea* n=1). Genetic distances between samples were computed with ANGSD version 0.937 (Korneliussen et al., 2014) using single-read resampling identity-by-state (IBS) algorithm, with the following filtering parameters: base call quality greater than Q25, mapping quality greater than 20, SNP p-value less than 1 x 10^−5^, a minimum minor allele frequency of 0.05, and sequenced in 75-80% of samples. Reads with multiple best hits and triallelic sites were discarded. These filters retained 7,238 SNPs for *M. cavernosa* and 16,147 SNPs for *S. siderea*. Using the *hclust* function in R version 4.0.3 (R Core Team, 2020), we created a hierarchical clustering tree to identify clones by qualitatively comparing clustering similarity to known technical replicates. One individual from each set of clones was retained for downstream analyses (sets of clones: *M. cavernosa* n=2, *S. siderea* n=5).

With clones removed, we again identified variant SNPs with the ANGSD parameters described above and identified probable admixture assignment with NGSadmix (Skotte et al., 2013), part of the ANGSD program suite. Individuals with greater than 25% assignment in an alternative lineage were considered hybrids. The IBS matrix produced an ordination (principal coordinate analysis, PCoA) using the function *capscale* from the R package *vegan* (Oksanen et al., 2020).

### Evaluation of differential gene expression

Raw TagSeq reads were first trimmed and cleaned using the custom script tagseq_clipper.pl (https://github.com/z0on/tag-based_RNAseq/blob/master/tagseq_clipper.pl). Because TagSeq focuses sequencing effort on the 3’ end of mRNA (Lohman et al., 2016), we added a 300-base buffer to the 3’ end of all annotated genes in the genome of *M. cavernosa* using the custom script add_3prime_buffer_to_gff.py (https://github.com/dgallery/FLKeys_trans2_adults/blob/main/add_3prime_buffer_to_gff.py). We then used the program featureCounts version 2.0.3 (Liao et al., 2014) to summarize the read counts. For *S. siderea* we used the custom scripts samcount_launch_bt2.pl (https://github.com/z0on/tag-based_RNAseq/blob/master/samcount_launch_bt2.pl) and expression_compiler.pl (https://github.com/z0on/tag-based_RNAseq/blob/master/expression_compiler.pl) using the annotated transcriptome (Davies et al., 2016) as reference for read mapping. Genes with a mean read count of less than five were removed from the data prior to downstream analyses, leaving 9479 genes for *M. cavernosa* and 7559 genes for *S. siderea*. Gene expression counts were normalized using variance stabilized transformation using the *vst* and *assay* functions in the R package *DESeq2* version 1.30.1 (Love et al., 2014).

To test whether differential gene expression was primarily driven by habitat or lineage affiliation, we first used the *capscale* function in the R package vegan (Oksanen et al., 2020) to obtain principal coordinates for both species. As the distance measure, we used Manhattan distance on the variance-stabilized counts (generated using function *vst* of the *DESeq2* package) averaged across all genes, corresponding to the average log_2_-fold difference in expression. We evaluated the number of significant principal components (*M. cavernosa*, n = 2; *S. siderea*, n = 3) using the R function *vegan::bstick* (Oksanen et al., 2020) and plotted the resulting principal coordinate analyses (PCoAs) colored by habitat and lineage. Using the *adonis2* function in the R package vegan (Oksanen et al., 2020), we performed a permutational multivariate analysis of variance with the distance matrices obtained from the *capscale* function with 1×10^6^ permutations and a set seed for each run. We evaluated the regression and residuals for habitat and lineage to determine which best explained the gene expression patterns in the data. We then used a likelihood ratio test (LRT) in *DESeq2* to analyze the effect of habitat and lineage using a reduced model approach.

We identified modules of highly correlated genes using a weighted gene co-expression network analysis (WGCNA) (Langfelder & Horvath, 2008). WGCNA analyzes the correlation between different genes and identifies clusters of genes that are expressed in similar ways blinded to the study design. Gene cluster identification can then be referenced against study design parameters (e.g., habitat, lineage) to identify trends in the data. We identified a soft threshold power of 8 (*M. cavernosa*) and 10 (*S. siderea*) and set the minimum module size to 30 genes for both species. We then merged similar gene modules, which resulted in a total of n = 6 modules for both species. We then compared overall differential gene expression within each module with DNA-based lineage membership and habitat location with heatmaps.

To analyze the predicting variables of gene expression within and between lineages and habitats, we used the R package *gradientForest* (Ellis et al., 2012) to analyze WGCNA module eigengene connectivity (kMEs). Gradient forest is a version of random forest that uses bootstrapped regression trees to determine the predictors of a response variable (in this case, gene expression) and captures the interactions between predictors (Ellis et al., 2012). Unlike classical generalized linear models in *DESeq2*, gradient forest can automatically choose important variables, account for their interactions without the need to specify them and without loss of power, detect linear as well as non-linear relationships, and perform cross-validations to measure the importance of the predictor variables.

To determine potential genes of interest, we identified WGCNA modules that were differentially expressed between habitats and/or lineages. Utilizing Gene Ontology with a Mann-Whitney U test (GO-MWU), we identified significant (p < 0.05) GO terms from each species (Wright et al., 2015, https://github.com/z0on/GO_MWU). We analyzed the resulting GO terms in three classes (molecular functions, cellular components, and biological processes) to examine potential differences in biological actions between habitats and lineages.

### Comparison of genetic divergence to differential gene expression

To determine if genes with high levels of genetic divergence also experience large levels of differential gene expression, we compared *F*_ST_ to patterns of differential gene expression. Because we lack an annotated genome for *S. siderea*, this analysis could only be done on *M. cavernosa*. *F*_ST_ was calculated in ANGSD version 0.937 (Korneliussen et al., 2014) using the Fumagalli method (Fumagalli et al., 2013, 2014). Variant SNP sites were identified with the following quality control parameters: base call quality greater than Q35, mapping quality greater than Q30, SNP p-value less than 1 x 10^−3^, a maximum heterozygosity frequency of 0.5, and sequenced in 80% of samples. Sites were then indexed, and the site allele frequency likelihoods were calculated for each subpopulation: Nearshore lineage, Offshore lineage, and Deep 1 lineage. Deep 2 lineage was excluded from this analysis because only one individual from this lineage was identified in the West transect. Two-dimensional site frequency spectra (2D-SFS) were made the following pairwise comparisons: Nearshore lineage vs. Offshore lineage, Nearshore lineage vs. Deep 1 lineage, and Offshore lineage vs. Deep 1 lineage. Per-site *F*_ST_ was calculated for each pairwise comparison using the annotated genome assigned to the gene in which the SNP occurred. In genes with multiple SNPs, *F*_ST_ was calculated by summing the alpha components and dividing by the sum of the alpha and beta components (Fumagalli et al., 2014). The SNP location was compared to the annotated genome to determine the gene in which each occurred. The SNP with the maximum minor allele frequency was retained in genes with multiple SNPs.

We then compared the *F*_ST_ of the top genes from each WGNCA module to see if there were differences in the frequency of *F*_ST_ values between modules, particularly those that showed differences between lineages. Each WGCNA module assigns a value of how much each gene belongs to that module, kME (correlation of the gene’s expression with the module’s eigengene). Using this value, we selected the top 100 genes representing each module and plotted quantiles of the logit-*F*_ST_ for these genes against logit-*F*_ST_ quantiles for 100 random genes from the genome to determine if any of the modules contained genes with higher *F*_ST_ than expected. The confidence limits for this comparison were determined by sampling 100 random genes repeatedly (100 times) and comparing their logit-*F*_ST_ quantiles.

## Results

### Population structure

We tested whether the West transect (this dataset) contained the four lineages described previously in the East transect (Rippe et al., 2021). Using NGSadmix (Skotte et al., 2013), we tested cluster assignment independently by combining the East transect (Rippe et al., 2021) and the West transect (this dataset) and with only the West transect for both *M. cavernosa* and *S. siderea*. Cluster assignments remained consistent, and we confirmed that both species have four primary lineages in this region. With the combined IBS matrix of the East and West transect, we created a principal coordinate analysis (PCoA) overlapping the East and West transects (Figure 1b, *M. cavernosa* and Figure 1d, *S. siderea*). We observed high fidelity between lineage assignments across both transects. These data show the robustness of 2bRAD with no batch effects between the two datasets despite them being processed entirely independently years apart. Finally, we compared the relative abundance of each lineage by depth for both transects (Figure 1c and 1e; *M. cavernosa* and *S. siderea*, respectively). These results confirm those of Rippe et al. (2021), which found two lineages specialized primarily to the deep environment and two primarily specialized to the shallow environments.

### Drivers of gene expression

We used three methods to determine whether cryptic lineage identity or environment is the primary driver of gene expression in *M. cavernosa* and *S. siderea*: gradient forest, *DESeq2*, and WGCNA. The gradient forest analysis results indicated that within *M. cavernosa*, nearshore and offshore habitats are the two strongest predictors of gene expression variation (Figure 2a). The Deep 1 lineage is a slightly stronger predictor of differential gene expression than the deep habitat (Figure 2a). In *S. siderea*, all three habitats are stronger predictors of gene expression variation than the lineages (Figure 2c). These data are supported by the *DESeq2* results, where the likelihood ratio test indicated out of 8546 genes analyzed for *M. cavernosa,* habitat predicted 879 differentially expressed genes, and lineage predicted 18 differentially expressed genes (Figure 2a inset). In 6610 genes analyzed in *S. siderea* habitat predicted 2586 differentially expressed genes, and lineage predicted 69 differentially expressed genes (Figure 2c inset). Furthermore, the WGCNA results show more significant results when samples are clustered by habitat rather than lineages (Supplementary Figure 1).

**Figure 2:**
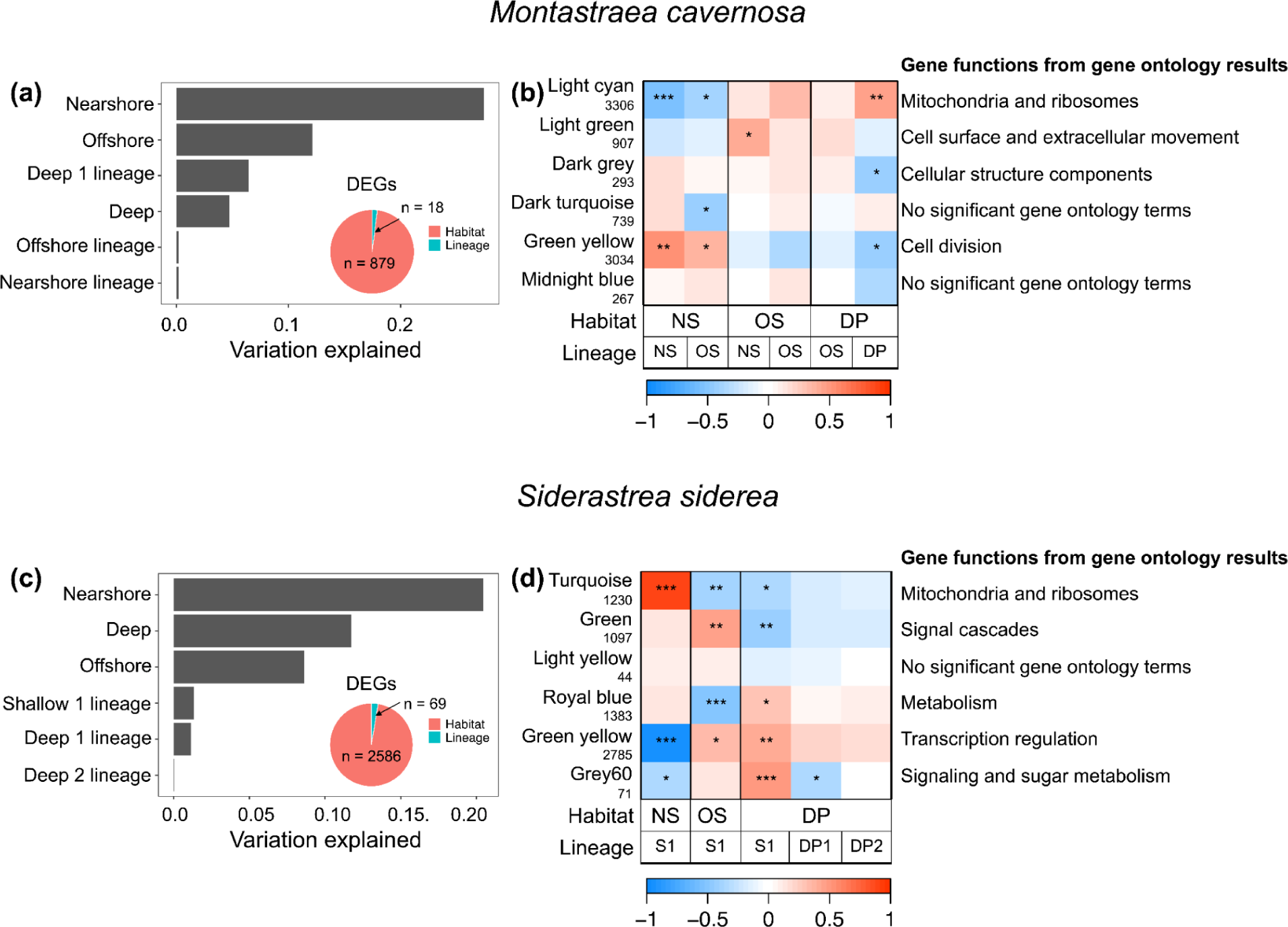
Lack of lineage-specific responses to habitat type at the gene expression network level. (a, c): Association between the identified gene network modules and predictors (habitat type or lineage), according to the gradient forest analysis accounting for possibly non-linear relationships and interactions and based on cross-validation. The length of each bar gives the proportion of variation predicted by a specific factor level. Insets are pie charts giving the number of differentially expressed genes (“DEGs”) with respect to either habitat or lineage, according to gene-by-gene generalized linear modeling in *DESeq2*. (b, d): Heatmaps showing results of linear regression for the same associations. “NS” - nearshore, “OS” - offshore, “DP” - deep. * = p ≤ 0.05, ** = p ≤ 0.01, *** = p ≤ 0.001. (a, b) *M. cavernosa*. (c,d) *S. siderea.* Gene ontology terms for the module are placed to the right of the heatmap.

Using WGCNA, a non-supervised method that clusters genes with similar expression profiles across samples, we identified six modules of coregulated genes in both *M. cavernosa* (Figure 2b) and *S. siderea* (Figure 2d). To quantify the importance of habitat and lineage in explaining the expression of these modules, we used the gradient forest approach, using lineage and habitat as predictors. Since gradient forest requires numerical predictors, these categorical variables were represented as several “dummy” quantitative variables (with values 1 or 0) corresponding to each category. In both species, the most important predictor was the “Nearshore” habitat type, explaining over 20% of the total variation (Fig. 2 a,c). Overall, habitat explained 43% of the variation in *M.cavernosa* and 41% of the variation in *S.siderea*, while lineage explained only 7% of the variation in *M.cavernos*a and 2.5% in *S.siderea*. Note that under the gradient forest approach, these numbers account for all possible lineage:habitat interactions and possible non-linear dependencies. The dominance of habitat in explaining the module eigengene expression was also visible when plotting a heatmap showing the slopes of simple linear regressions of module eigengene expression against the same dummy variables (Fig. 2 b,d).

We also analyzed components of the variance-stabilized gene expression counts using the *capscale* function with Manhattan distances (corresponding to the average log-fold difference across all genes). The resultant PCoAs (Supplementary Figure 2) were colored by habitat and lineage. We evaluated the statistical significance of each predictor variable (habitat and lineage) with the permutational multivariate analysis function, *adonis2*. Results indicate an R^2^ of 0.50 (*M. cavernosa*, p = 1×10^−6^) and 0.45 (*S. siderea*, p = 1×10^−6^) when habitat is the primary predictor variable, and a regression of 0.28 (*M. cavernosa*, p = 2.0 x 10^−6^) and 0.10 (*S. siderea*, p = 0.01) when lineage is the primary predictor variable. The aggregate of these results strongly supports that environment, rather than genetics, is the primary determinant for gene expression differentiation in *M. cavernosa* and *S. siderea*.

We then analyzed each WGNCA module for the two species for gene ontology terms using GO-MWU. We summarized the most prominent associated gene functions for each module in each species (Figure 2b, *M. cavernosa,* and Figure 2d, *S. siderea*). Full lists of significantly associated terms are available in the supplementary figures (Supplementary Figures 3 and 4). We expected to find modules were regulated primarily by lineage; however, instead, we found most modules were similar within a habitat regardless of an individual’s lineage membership. Instead, we found several WGCNA modules showing differential expression across habitats. differential gene expression in different habitats. Notably, for *M. cavernosa*, the lightcyan and lightgreen modules in both Nearshore and Offshore genetic lineages are downregulated in the nearshore habitat but upregulated in the offshore habitat (Figure 2b, Supplementary Figure 3). These modules are associated with mitochondria and ribosomes, and cell surface and extracellular movement, respectively. In the darkgrey module, where lineage seems to predict gene expression rather than habitat, gene ontology terms are primarily cellular structure components (Figure 2b, Supplementary Figure 3).

*S. siderea* also has opposing responses in different habitats despite belonging to the same lineage, as demonstrated in the heatmap (Figure 2d). Because the majority of the samples for this species belong to the Shallow 1 lineage, we focus only on this lineage. This pattern is seen in the turquoise, green, royalblue, greenyellow, and grey60 modules. These modules correspond to cellular growth, signal cascades, metabolism, transcription regulation, and signaling and sugar metabolism, respectively (Figure 2d, Supplementary Figures 4). There are no WGNCA modules in *S. siderea* with the same differential gene expression in all three habitats for this lineage.

### Genetic divergence of genes in coregulated modules (*M. cavernosa*)

We expected that modules with lineage-specific expression in the same habitat, i.e., darkgrey and midnightblue, would have higher *F*_ST_ values compared to modules without lineage-specific expression, i.e., lightcyan and lightgreen, or compared to a random selection of genes from the genome. However, none of the modules appeared to be enriched with high-*F*_ST_ genes (Figure 3). If they were, their quantile-quantile (q-q) plots would deviate upwards from the diagonal in Figure 3. Instead, they don’t depart from the random expectation (comparison between two randomly chosen groups of genes).

**Figure 3:**
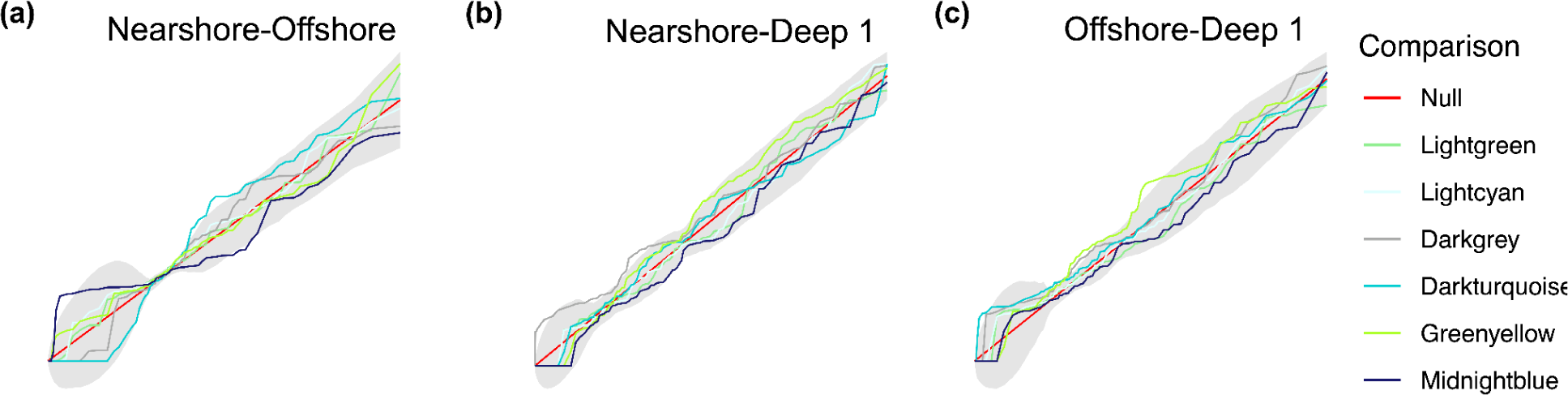
No evidence of accelerated genetic divergence in any of the gene expression modules in *M. cavernosa*. Quantile (Q-Q) plots of logit-*F*_ST_ of the top 100 genes belonging to each WGCNA module compared to 100 randomly chosen genes from the genome (Null, colored red). The gray area around the diagonal is 1.96*SD of the contrast between two random draws of 100 genes. (a) *F*_ST_ comparison between Nearshore lineage and Offshore lineage. (b) *F*_ST_ comparison between Nearshore lineage and Deep 1 lineage. (c) *F*_ST_ comparison between Offshore lineage and Deep 1 lineage. None of the graphs shows a significant departure upward from the diagonal at the top-left corner, which would indicate an excess of large logit-Fst values.

## Discussion

### Population Structure

Our results indicate that this dataset includes the same lineages originally described in Rippe *et al*. (2021), albeit at different frequencies than previously observed. Specifically, *M. cavernosa* only had one individual in the Deep 2 lineage, and *S. siderea* had no individuals in the Shallow 2 lineage. Interestingly, in *M. cavernosa,* the Deep 2 lineage also had the lowest proportion of individuals in the East transect, and only 2 of those individuals were identified as juvenile corals less than 3 cm in diameter. This reduction of the Deep 2 lineage may be due to recruitment stochasticity or lower relative fitness compared to the Deep 1 lineages (e.g., natural selection). In the latter case, it is possible that the Deep 2 lineage is adapted to even deeper waters. Furthermore, the West transect samples were collected prior to the 2015 bleaching event, whereas the East transect samples were collected in 2018. This may indicate that the Deep 2 lineage has reduced heat tolerance compared to the Deep 1 lineage, which would result in a reduced abundance of this lineage across the transect in the 2018 sampling. Evaluating each lineage’s bleaching and recovery tolerance for both species will be vital for successful future restoration efforts.

The lack of the Shallow 2 lineage of *S. siderea* in the West transect may indicate low recruitment within this lineage in the Florida Keys, whereas the high level of the Shallow 1 lineage could be attributed to relatively high recruitment. Two other hypotheses may explain these data: demographic stochasticity and natural selection. The 2015 bleaching event affected *S. siderea,* causing many colonies to perish (Smith et al., 2019) and may indicate that other lineages did not have the heat tolerance required to survive this event. Again, a study of the relative heat tolerance of all lineages is required to test this hypothesis. Another hypothesis for this loss of the Shallow 2 lineage and reductions of the Deep 1 and Deep 2 lineages is that the Shallow 1 lineage is expanding in the Florida Keys, displacing all other lineages. Indeed, the majority of juvenile *S. sidere*a across all three habitats (even in the deep) belong to this lineage (Rippe *et al*. 2021).

### Shared acclimatization pathways between lineages

Previous research has shown that difference between genets is the primary driver of differential gene expression in corals (Dixon et al., 2015; Meyer et al., 2011). In two reciprocal transplantation experiments, one in Caribbean *Porites astreoides* (Kenkel & Matz, 2016) and another in Indo-Pacific *Acropora millepora* (Dixon et al., 2018), between-genet difference accounted for over 60% of total gene expression variation, despite the fact that the fragments have been planted in very different habitats for weeks or even months. We, therefore, hypothesized that gene expression patterns would follow coral genetics more than an environment: that they would be more similar within lineages rather than within habitats. However, PERMANOVA, gene-by-gene generalized linear models, gradient forest, and WGCNA analyses all indicate that differential gene expression is predicted by environmental conditions much more than by cryptic lineage affiliation (Figure 2, Supplementary Figures 1-2). Despite the measurable depth specialization of each lineage (Figure 1c, e), their gene expression profiles converge when found in the same habitat (Figure 2), indicating shared gene regulatory mechanisms involved in acclimatization. It is possible that this similarity in plastic responses between lineages within a species is due, at least in part, to the ongoing partial introgression between cryptic lineages (Rippe et al., 2021). However, because each lineage retains enough gene expression plasticity to survive in different environments, it begs the question of how these lineages formed initially since being locally adapted to a specific environment is no longer supported as the primary reason. These results surprisingly indicate that lineage-specific gene expression is not the primary driver of the local adaptation in our study species. If so, adaptations to the deep and shallow environments must have occurred via other mechanisms, such as modification of the genes’ sequences rather than expression levels or within a small number of genes that were too few to form a module in our WGCNA analysis. It is also possible that the ecological specialization of lineages is driven primarily by their evolved symbiotic associations with divergent Symbiodiniaceae and/or bacterial communities instead of adjustments to the coral host’s own physiology.

The hypothesis that *M. cavernosa* and *S. siderea* may be utilizing their endosymbionts and microbiomes to adapt to their local environment is particularly interesting. One study observed that coral transcriptomes within a single clone of *Orbicella faveolata* were predicted by their symbiont’s clade rather than the environmental variation (DeSalvo et al., 2010). In *Pocillopora damicornis,* the symbionts and microbes were found to be locally adapted to habitats and potentially provide the coral with the adaptive variation necessary to survive in various environments (van Oppen et al., 2018). While *M. cavernosa* is usually associated with the *Cladocopium* genus of zooxanthellae (Serrano et al., 2014), that genus may include a large amount of diversity within it. To explore this issue, we need a proper population genomics analysis to accurately compute genome-wide genetic distances between symbiont clones. Reciprocal transplant and common garden experiments monitoring the shifts in zooxanthellae composition, microbial community composition, and their functional contributions to host fitness could provide the necessary samples.

### Lack of correlation between *F*_ST_ and differential gene expression within *M. cavernosa*

We identified two modules of coregulated genes that varied by lineage rather than by habitat: darkgrey (n = 293) and midnightblue (n = 267). If the divergence of their expression was helped by natural selection (during adaptation to a specific habitat), one would expect these modules to be enriched with genes with elevated *F*_ST_ compared to other genes in the genome. However, this does not seem to be the case (Figure 3). In the eastern oyster, *Crassostrea virginica*, genes with diverging gene expression values were found to be correlated with higher-than-average *F*_ST_ values (Johnson et al., 2021). However, in *Salvelinus alpinus*, the Arctic char, similar results to ours were found, where differential gene expression did not correlate with higher than average *F*_ST_ values (Jacobs et al., 2020). Jacobs et al. (2020) hypothesized that parallel evolution caused gene expression to converge due to environmental effects. In our case, it is possible that small modules with lineage-specific expression unite genes that experienced neutral expression divergence between lineages and are not, in fact, related to their ecological specialization.

## Conclusion

*Montastraea cavernosa* and *Siderastrea siderea* in the Florida Keys each comprise four distinct genetic lineages, unevenly distributed by the nearshore-offshore gradient and especially by depth. While regulatory evolution seems like a plausible mechanism to produce such ecological specialization, we did not find evidence for it. The remaining possibilities include adjustments to genes’ sequence affecting the proteins’ function rather than their expression level, adjustment of expression of a small number of genes (less than 30) that would not be detectable as a gene network module, or change in composition of algal symbionts or microbiome rather than evolution of the coral host.

## Acknowledgments

Funding for this project was provided by the National Science Foundation grant OCE-1737312 to M.V.M, and all collections were authorized under the Florida Keys National Marine Sanctuaries permits 2014-072A and 2015-071. We wish to thank Eric Bartels and Mote’s Elizabeth Moore International Center for Coral Reef Research & Restoration for invaluable field assistance and resources. The bioinformatics analysis was accomplished using computational resources provided by the Texas Advanced Computer Center. We also wish to thank Hayley Bedwell for her assistance in the field, Coral Loockerman for her assistance with laboratory work, and Evelyn Abbott, Kristina Black, and Carly Scott for their assistance with data analysis.

## Data Accessibility

Raw 2bRAD sequence data from this study can be accessed under the NCBI BioProject Accession PRJNA813607. The *Montastraea cavernosa* reference genome is available on the Matz Lab website (https://matzlab.weebly.com/data--code.html). Bioinformatic procedures associated with the de novo 2bRAD methodology (https://github.com/z0on/2bRAD_denovo), bioinformatic procedures associated with the TAG-seq methodology (https://github.com/z0on/tag-based_RNAseq), and all other data analysis procedures used in this study (https://github.com/dgallery/FLKeys_trans2_adults) are available in the specified GitHub repositories.

## Author contributions

M.M. conceived and designed this study. M.M. and J.R. collected tissue samples in the field, and J.R. performed 2bRAD and TAG-seq library preparations. D.G., J.R., and M.M. conducted the bioinformatic and data analysis. D.G. and M.M. prepared the manuscript with all authors contributing to its final form.

## Notes

### Competing Interest Statement

The authors have declared no competing interest.

