## Supplementary Figures 1-4 for "Decrypting corals: Does regulatory evolution underlie environmental specialization of coral cryptic lineages?"

Supplementary Materials

Figures:

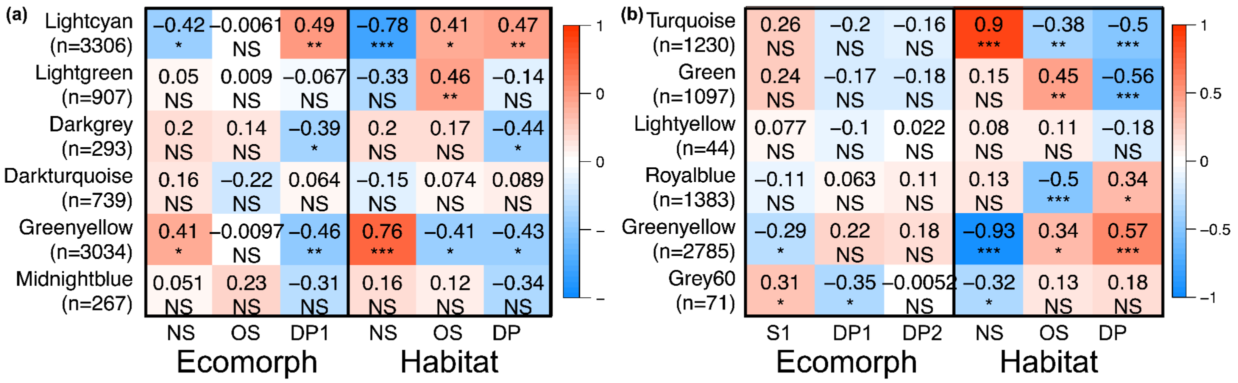


Supplementary Figure 1: WGNCA modules separated by lineage and habitat for *M. cavernosa* (a) and *S. siderea* (b).

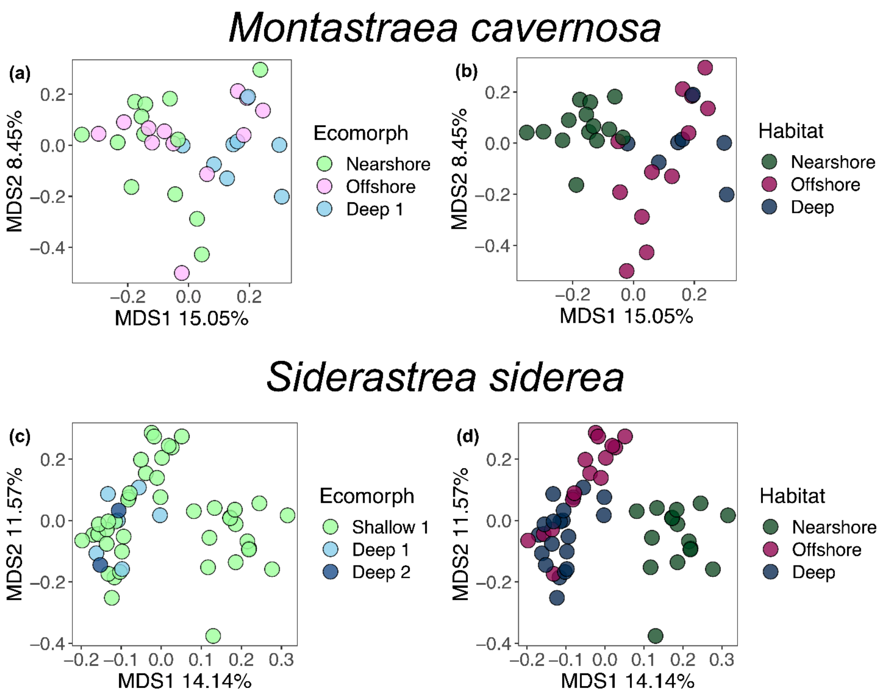


Supplementary Figure 2: PCoA of variance stabilized gene expression counts for *M. cavernosa* (a, b) and *S. siderea* (c,d), colored by habitat (a, c) and lineage (b, d).

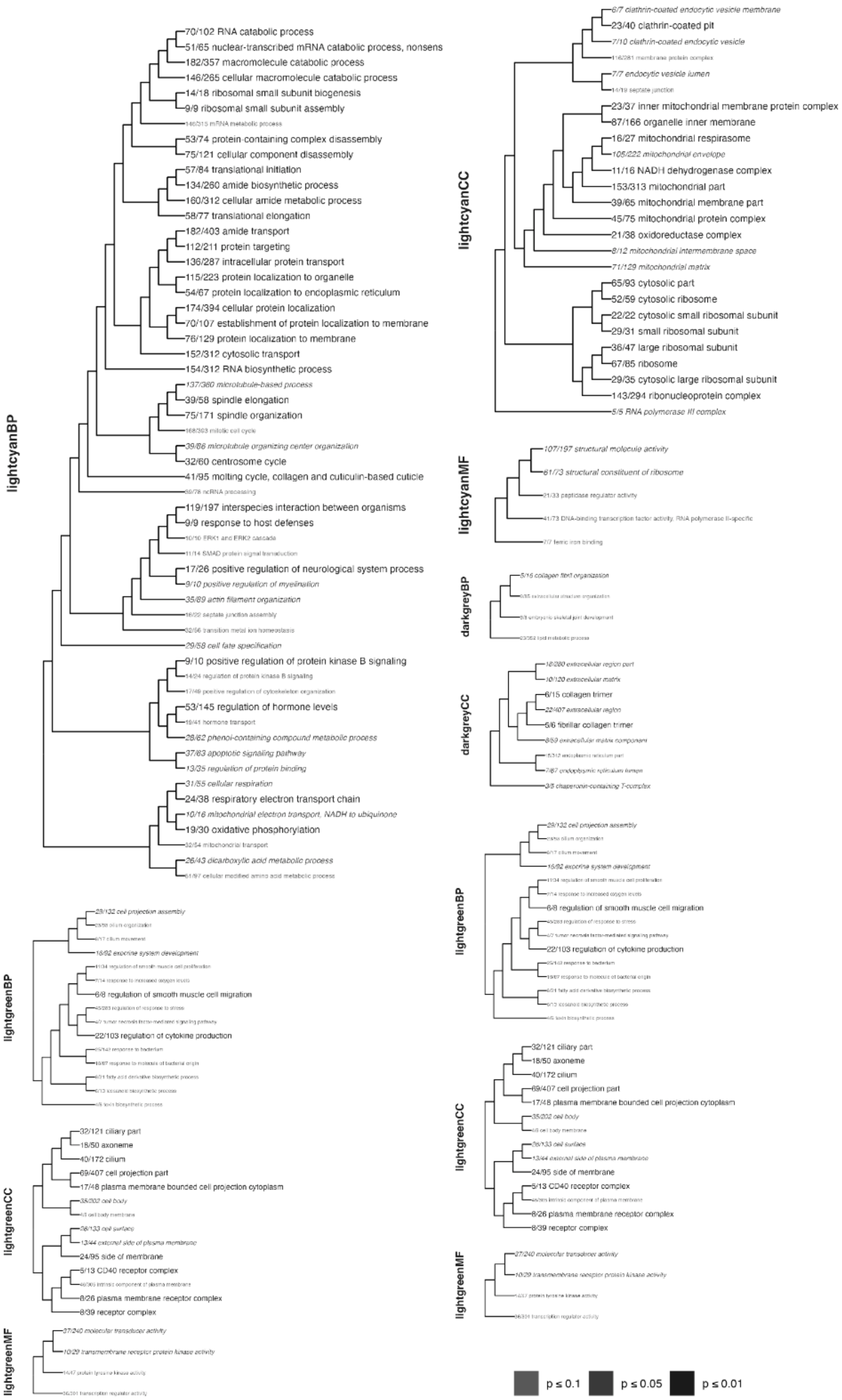

lightgreenBP
29/132 cell projection assembly
6/17 cilium movement
16/92 exocrine system development
6/8 regulation of smooth muscle cell migration
22/103 regulation of cytokine production
lightgreenCC
32/121 ciliary part
18/50 axoneme
40/172 cilium
69/407 cell projection part
17/48 plasma membrane bounded cell projection cytoplasm
35/202 cell body
28/133 cell surface
13/44 external side of plasma membrane
24/95 side of membrane
5/13 CD40 receptor complex
8/26 plasma membrane receptor complex
8/39 receptor complex
lightgreenMF
27/240 molecular transducer activity
10/29 transmembrane receptor protein kinase activity

Supplementary Figure 3: Gene ontology terms for *M. cavernosa*.

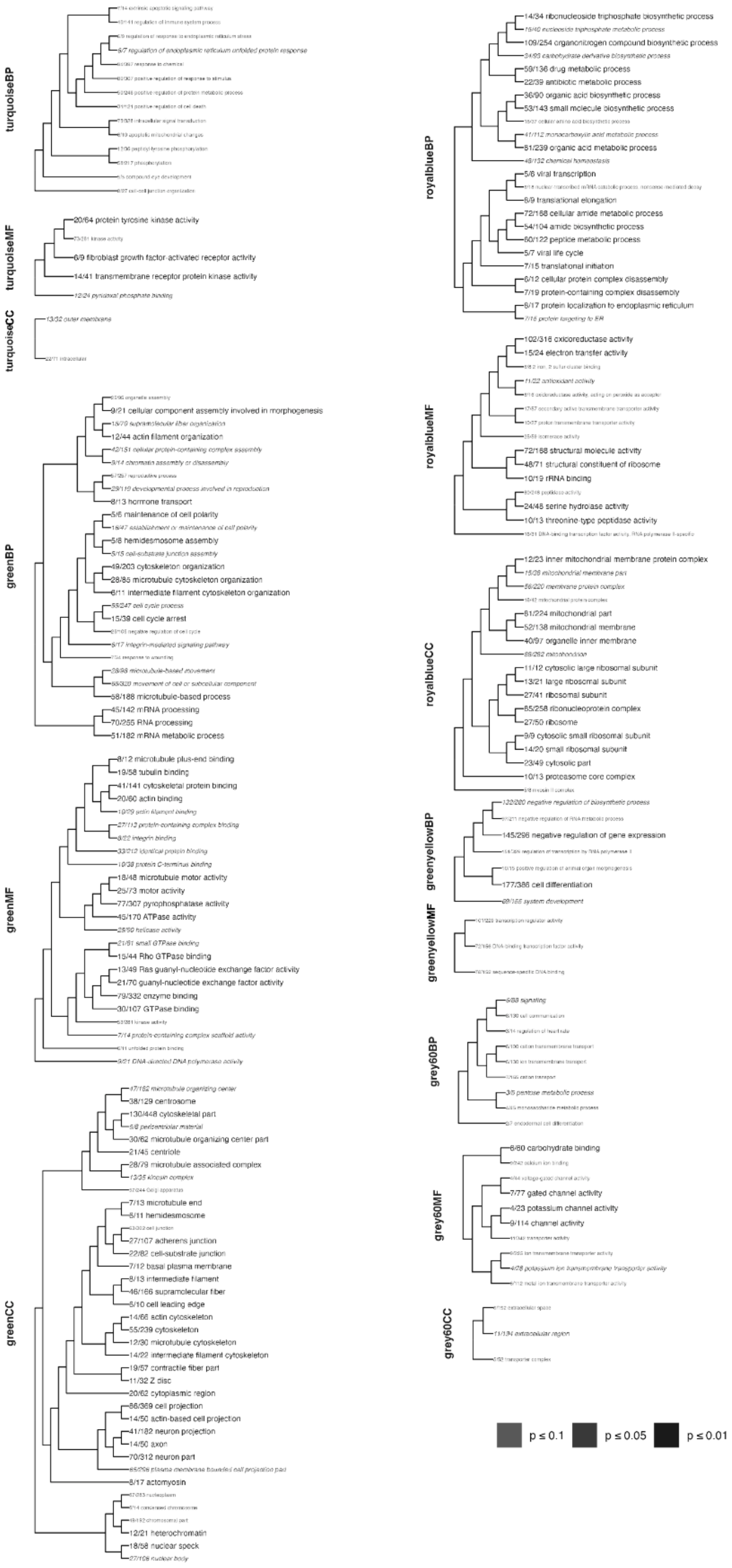

turquoiseBP
9/7 regulation of endoplasmic reticulum unfolded protein response
turquoiseMF
20/64 protein tyrosine kinase activity
6/9 fibroblast growth factor-activated receptor activity
14/41 transmembrane receptor protein kinase activity
12/24 pyridoxal phosphate binding
turquoiseCC
13/32 outer membrane
greenBP
9/21 cellular component assembly involved in morphogenesis
18/79 supramolecular fiber organization
12/44 actin filament organization
42/151 cellular protein-containing complex assembly
9/14 chromatin assembly or disassembly
29/119 developmental process involved in reproduction
8/13 hormone transport
5/6 maintenance of cell polarity
16/47 establishment or maintenance of cell polarity
5/8 hemidesmosome assembly
5/15 cell-substrate junction assembly
49/203 cytoskeleton organization
28/85 microtubule cytoskeleton organization
6/11 intermediate filament cytoskeleton organization
55/247 cell cycle process
15/39 cell cycle arrest
6/17 integrin-mediated signaling pathway
28/98 microtubule-based movement
85/320 movement of cell or subcellular component
58/188 microtubule-based process
45/142 mRNA processing
70/255 RNA processing
51/182 mRNA metabolic process
greenMF
8/12 microtubule plus-end binding
19/58 tubulin binding
41/141 cytoskeletal protein binding
20/60 actin binding
10/29 actin filament binding
27/112 protein-containing complex binding
8/22 integrin binding
33/212 identical protein binding
10/38 protein C-terminus binding
18/48 microtubule motor activity
25/73 motor activity
77/307 pyrophosphatase activity
45/170 ATPase activity
28/90 helicase activity
21/61 small GTPase binding
15/44 Rho GTPase binding
13/49 Ras guanyl-nucleotide exchange factor activity
21/70 guanyl-nucleotide exchange factor activity
79/332 enzyme binding
30/107 GTPase binding
7/14 protein-containing complex scaffold activity
9/21 DNA-directed DNA polymerase activity
greenCC
47/182 microtubule organizing center
38/129 centrosome
130/448 cytoskeletal part
6/8 pericentriolar material
30/62 microtubule organizing center part
21/45 centriole
28/79 microtubule associated complex
13/35 kinesin complex
7/13 microtubule end
6/11 hemidesmosome
27/107 adherens junction
22/82 cell-substrate junction
7/12 basal plasma membrane
8/13 intermediate filament
46/166 supramolecular fiber
5/10 cell leading edge
14/66 actin cytoskeleton
55/239 cytoskeleton
12/30 microtubule cytoskeleton
14/22 intermediate filament cytoskeleton
19/57 contractile fiber part
11/32 Z disc
20/62 cytoplasmic region
86/369 cell projection
14/50 actin-based cell projection
41/182 neuron projection
14/50 axon
70/312 neuron part
85/296 plasma membrane bounded cell projection part
8/17 actomyosin
12/21 heterochromatin
18/58 nuclear speck
27/108 nuclear body
royalblueBP
14/34 ribonucleoside triphosphate biosynthetic process
15/40 nucleoside triphosphate metabolic process
109/254 organonitrogen compound biosynthetic process
34/95 carbohydrate derivative biosynthetic process
59/136 drug metabolic process
22/39 antibiotic metabolic process
36/90 organic acid biosynthetic process
53/143 small molecule biosynthetic process
41/112 monocarboxylic acid metabolic process
81/239 organic acid metabolic process
48/132 chemical homeostasis
5/8 viral transcription
8/9 translational elongation
72/168 cellular amide metabolic process
54/104 amide biosynthetic process
60/122 peptide metabolic process
5/7 viral life cycle
7/15 translational initiation
6/12 cellular protein complex disassembly
7/19 protein-containing complex disassembly
8/17 protein localization to endoplasmic reticulum
7/16 protein targeting to ER
royalblueMF
102/316 oxidoreductase activity
15/24 electron transfer activity
11/22 antioxidant activity
72/168 structural molecule activity
48/71 structural constituent of ribosome
10/19 rRNA binding
24/48 serine hydrolase activity
10/13 threonine-type peptidase activity
royalblueCC
12/23 inner mitochondrial membrane protein complex
15/26 mitochondrial membrane part
56/220 membrane protein complex
61/224 mitochondrial part
52/138 mitochondrial membrane
40/97 organelle inner membrane
86/262 mitochondrion
11/12 cytosolic large ribosomal subunit
13/21 large ribosomal subunit
27/41 ribosomal subunit
65/258 ribonucleoprotein complex
27/50 ribosome
9/9 cytosolic small ribosomal subunit
14/20 small ribosomal subunit
23/49 cytosolic part
10/13 proteasome core complex
greenyellowBP
132/280 negative regulation of biosynthetic process
145/296 negative regulation of gene expression
177/386 cell differentiation
89/165 system development
greenyellowMF
grey60BP
6/83 signaling
3/5 pentose metabolic process
grey60MF
6/80 carbohydrate binding
7/77 gated channel activity
4/23 potassium channel activity
9/114 channel activity
4/28 potassium ion transmembrane transporter activity
grey60CC
11/134 extracellular region
p ≤ 0.1
p ≤ 0.05
p ≤ 0.01

Supplementary Figure 4: Gene ontology terms for *S. siderea.*
